# Hidden multivalency in phosphatase recruitment by a disordered AKAP scaffold

**DOI:** 10.1101/2021.05.23.445342

**Authors:** Matthew Watson, Teresa B. Almeida, Arundhati Ray, Christina Hanack, Rory Elston, Joan Btesh, Peter A. McNaughton, Katherine Stott

## Abstract

Disordered scaffold proteins provide multivalent landing pads that, *via* a series of embedded Short Linear Motifs (SLiMs), bring together the components of a complex to orchestrate precise spatial and temporal regulation of cellular processes. One such protein is AKAP5, which contains SLiMs that anchor PKA and Calcineurin, and recruit substrate (the TRPV1 receptor). Calcineurin is anchored to AKAP5 by a well-characterised PxIxIT SLiM. Here we show, using a combination of biochemical and biophysical approaches, that the Calcineurin PxIxIT-binding groove also recognises several hitherto unknown lower-affinity SLiMs in addition to the PxIxIT motif. We demonstrate that the assembly is in reality a complex system with conserved SLiMs spanning a wide affinity range. The capture is analogous to that seen for many DNA-binding proteins that have a weak non-specific affinity for DNA outside the canonical binding site, but different in that it involves (i) two proteins, and (ii) hydrophobic rather than electrostatic interactions. It is also compatible with the requirement for both stable anchoring of the enzyme and responsive downstream signalling. We conclude that the AKAP5 C-terminus is enriched in lower-affinity/mini-SLiMs that, together with the canonical SLiM, maintain a structurally disordered but tightly regulated signalosome.

## Introduction

Scaffolding proteins are central to strict spatial and temporal control of cellular processes (Scott and Pawson, 2009; Pan *et al*., 2012). By increasing local concentrations, they enhance specificity and reaction rates of otherwise promiscuous enzymes. A-kinase anchoring proteins (AKAPs) are a family of intrinsically-disordered proteins (IDPs) that were originally characterized by their ability to scaffold Protein Kinase A (PKA) (Theurkauf and Vallee, 1982; Lohmann *et al*., 1984), but have since been shown to assemble components into multifunctional ‘signalosomes’ that can be targeted to specific receptors or sites (Wong and Scott, 2004; Torres-Quesada, Mayrhofer and Stefan, 2017). Currently there are 14 annotated AKAPs in humans (AKAP1 to AKAP14).

AKAPs are typical of many IDPs in that they interact with their partners *via* Short Linear Motifs (SLiMs) (Davey *et al*., 2012; Roey *et al*., 2014). The presence of multiple different SLiMs in AKAPs enables the combinatorial assembly of a dynamic multi-protein hub. Much of our current understanding of disordered signalling complexes is based on the structures of folded partners bound to their canonical peptide SLiMs. The rest of the IDP is either not resolved or is removed to facilitate structural studies. In many cases, the regions between SLiMs are assumed to be inert linkers, simply providing a length-dependent tuning of effective concentration (although predictions from physical models are not always reproduced experimentally, suggesting the reality is more complicated (Charlotte, Jendroszek and Kjaergaard, 2019)). The context in which IDP(R) partner interactions take place can also play an important role (Bugge *et al*., 2020). The flanking sequences around SLiMs, although disordered even in the complex, often contribute to binding (Liu *et al*., 2010). Moreover, there can be complex interplay between SLiMs (Davey *et al*., 2012). For example, clustering or overlapping SLiMs can modulate interactions through cooperativity or competition. Further, multiple low-affinity SLiMs can cooperate to increase the binding affinity. The cooperativity can take different forms: e.g. allovalency (where multiple epitopes along the ligand bind a single site on a partner (Klein, Pawson and Tyers, 2003)), or ‘fuzziness’ (where multiple interchangeable interaction sites exist on both the ligand and partner (Tompa and Fuxreiter, 2008)).

Here we investigate the interaction of the ubiquitous phosphatase Calcineurin (also known as PP2B) with the C-terminal region of the scaffolding protein AKAP5 (previously known as AKAP79 in humans and AKAP150 in mice). Calcineurin couples changes in intracellular calcium concentration to the phosphorylation state of several important substrates (Creamer, 2020). One example is the NFAT (Nuclear Factor of Activated T cells) family of transcription factors, which are dephosphorylated by Calcineurin, causing their translocation to the nucleus and stimulation of transcriptional programmes (Li, Rao and Hogan, 2011). Like other phosphatases, Calcineurin dephosphorylates sites with little sequence similarity (Donella-Deana *et al*., 1994; Li *et al*., 2013), and primarily recognises substrates or their scaffolding proteins through the interaction with SLiMs, such as the PRIEIT sequence in NFATc1 (Aramburu *et al*., 1998; Garcia-Cozar *et al*., 1998). PRIEIT is a canonical Calcineurin recognition motif obeying the consensus sequence PxIxIT, a so-called ‘PxIxIT’ SLiM. This interaction has a 25 µM affinity, which is within the optimal window for its signalling function (Li *et al*., 2007). AKAP5 also has a PxIxIT SLiM with which it engages Calcineurin, albeit one with a higher affinity – PIAIIIT – that is suited to anchoring (0.4 µM), while not so tight that downstream signalling is inhibited (Li *et al*., 2012).

In this study, we examine the AKAP5/Calcineurin interaction from the perspective of the IDP scaffold. Using Surface Plasmon Resonance (SPR), isothermal titration calorimetry (ITC) and NMR spectroscopy, we show that binding is far more extensive than the PxIxIT alone and we identify additional regions that recognise the same PxIxIT-binding groove of Calcineurin with µM - mM affinity. We hypothesise that these additional low-affinity sites are populated at the high concentrations of Calcineurin found in the cell, and that they help to capture and maintain the high-affinity anchored Calcineurin. Moreover, they provide a separate ‘labile but localised’ pool of rapidly dissociating and rebinding Calcineurin, which could be intercepted by proteins such as NFAT that compete for Calcineurin in order to enact optimal downstream signalling.

## Results

### The AKAP5c scaffold is highly disordered and monomeric

AKAP5 can loosely be divided into three regions (Fig. 1a). Here, we have chosen to focus on the C-terminal region (300-427; AKAP5c), which we define as the region of homology across various species (Supp. Fig. S1) following a large insertion of unknown function in the rodent orthologues. The sequence contains three known SLiMs. Two bind enzymes (Calcineurin (Dell’Acqua *et al*., 2002; Oliveria, Dell’Acqua and Sather, 2007) or PKA (Newlon *et al*., 2001)) and one binds substrates (TRPV1/4 (Zhang, Li and McNaughton, 2008; Btesh *et al*., 2013; Mack and Fischer, 2017)). The enzyme-bound SLiMs adopt defined secondary structures in the complexes: a β-strand in the case of Calcineurin (Li *et al*., 2012) and an α-helix in the case of PKA (Newlon *et al*., 2001). There is no structural information on the SLiM reported to bind TRPV1/4 (Btesh *et al*., 2013; Mack and Fischer, 2017).

**Figure 1:**
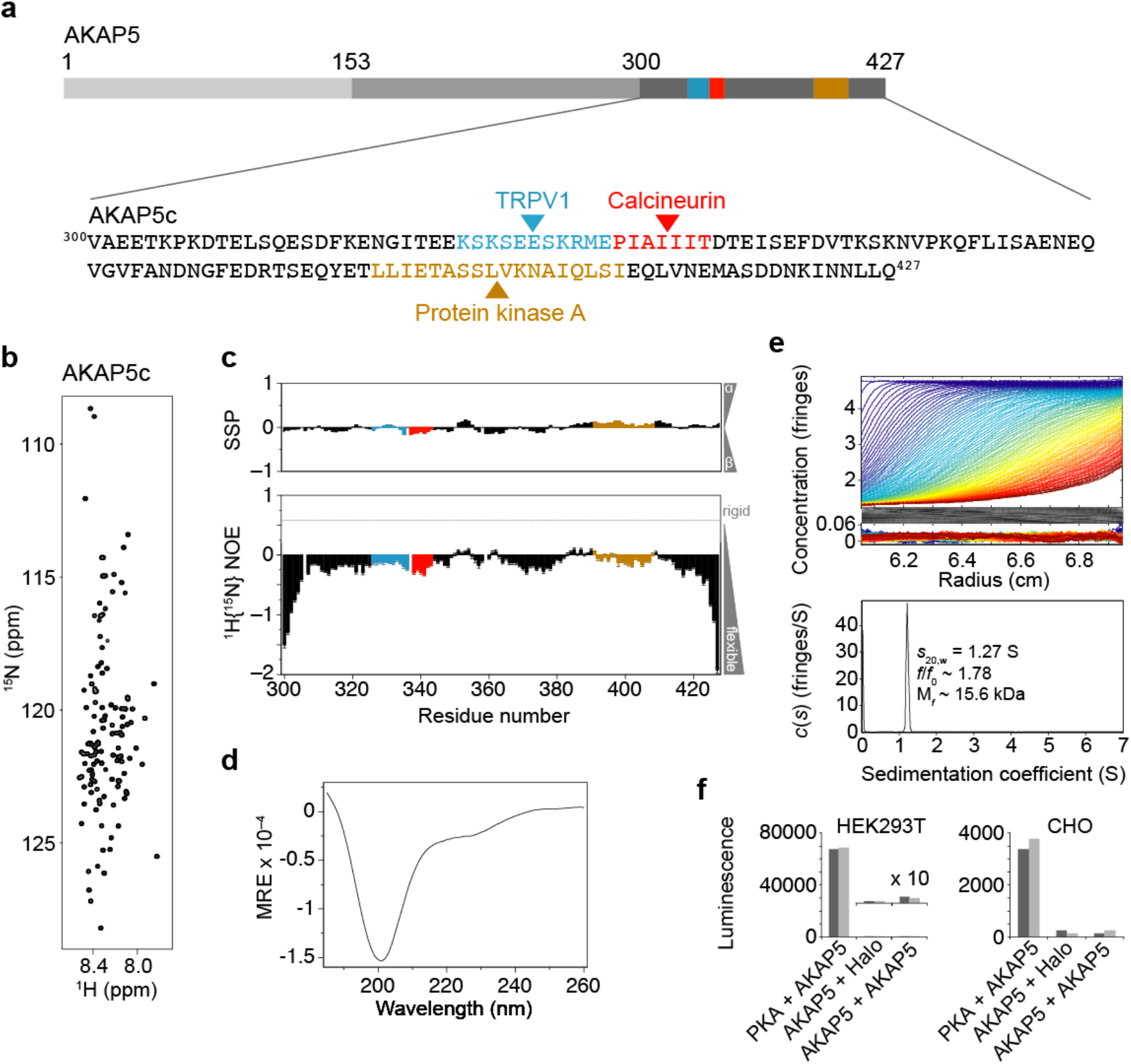
Structure, hydrodynamic properties and oligomeric state of AKAP5_300-427_ (AKAP5c). (**a**) Schematic of AKAP5: the N-terminal region (1-153) is involved in membrane attachment; the middle region (154-299) is recognised by scaffolding proteins of the Membrane associated Guanylate Kinase family; and the C-terminal region (300-427, shown expanded) contains three known SLiMs that bind Calcineurin, PKA and substrates e.g. TRPV1. (**b**) ^15^N HSQC of AKAP5c. (**c**) Secondary structure propensity (SSP) and Heteronuclear NOE values (colouring of SLiMs as **a**). (**d**) Far-UV CD spectrum. (**e**) Sedimentation velocity continuous *c*(*s*) distribution. (**f**) Live HEK- and CHO-cell split-luciferase assays (NanoBiT®, Promega). AKAP5 = full-length protein, PKA = regulatory PKA subunit (R2A) (positive control), Halo = HaloTag® (negative control). The order of listing indicates the orientation of the split luciferase: <protein>-LgBiT + <protein>-SmBiT.

To fully understand the role of disorder in the assembly of signalling complexes by scaffolding proteins, it is important to understand the level and type of disorder present. The term “intrinsic disorder” (Dunker *et al*., 2013) is used to describe all proteins that populate transient structures through to those that approach true random coils (i.e. those with no structural preferences and uniformly fast backbone dynamics). Two spectroscopic techniques were deployed to assess and delineate regions of nascent structure and to probe the backbone dynamics in AKAP5c: nuclear magnetic resonance (NMR) and circular dichroism (CD). The ^15^N-HSQC spectrum (Fig. 1b) displays the low ^1^H^N^ chemical-shift dispersion (0.7 ppm) and narrow line widths typical of disordered proteins. A near complete assignment of the backbone H, C, N and C^β^ chemical shifts was possible (Supp. Fig. S2), from which the secondary structure propensity (SSP (Marsh *et al*., 2006)) could be calculated (Fig. 1c). SSP scores in AKAP5c are close to zero with alternating stretches of weak α- or β-propensities. For comparison, SSP scores are ca. +1 or –1 in stable α- and β-structures, respectively. It is interesting to note that the propensities of the Calcineurin- and PKA-binding SLiMs (red and yellow, respectively, in Fig. 1c) reflect the stable structures adopted in the bound states (SSP_mean_ = –0.116 ± 0.038 and +0.079 ± 0.037, i.e. weak β and α, respectively). Backbone dynamics were probed by {^1^H}^15^N heteronuclear NOE (HNOE; Fig. 1c), which reveals motions on a time-scale faster than overall tumbling (HNOE < ca. +0.6). HNOEs were fairly uniform throughout AKAP5c (disregarding the highly mobile N- and C-termini). The average value was –0.20 ± 0.29, with no single residue exhibiting a value > +0.1, and all SLiMs falling within one standard deviation of the mean, consistent with a high level of disorder. CD spectroscopy also showed AKAP5c to be predominantly disordered (Fig. 1d): the signature large negative ellipticity at 200 nm was dominant, and although there were indications of some secondary structure (as evidenced by the weak shoulder at 222 nm), this is likely to be transiently populated given the results from NMR.

We next investigated the hydrodynamic properties of AKAP5c by sedimentation velocity analytical ultracentrifugation (SV AUC, Fig. 1e). AKAP5c sedimented slowly (*s*_20,w_ = 1.27) due to hydrodynamic friction resulting from its extended nature (frictional ratio *f*/*f*_0_ ∼ 1.78). The fitted mass (∼ 15.6 kDa) was close to the calculated monomer mass (14.5 kDa). The overall picture of AKAP5c is therefore one of a highly dynamic and monomeric chain. However, there are reports that AKAP5 can dimerise (Gold *et al*., 2011; Patel *et al*., 2017). Of particular relevance to this study is the observation of K333-K333 and K328-K328 intermolecular crosslinks (Gold *et al*., 2011), which are in the TRPV1/4 interaction SLiM immediately N-terminal to the Calcineurin binding site. Conversely, Nygren *et al*. (Nygren *et al*., 2017) concluded that the AKAP5/Calcineurin complex is monomeric by SEC-MALS. Given AKAP5 forms complex assemblies within the cell, we investigated whether full length AKAP5 dimerises in a cellular context (HEK293T and CHO) with a highly sensitive live-cell luminescence assay (NanoBiT®, Promega) that uses a split-luciferase reporter engineered to give reduced background. A strong complementation signal was seen between PKA and AKAP5 as expected for a direct PKA-AKAP interaction, however, AKAP5 did not give a self-complementation signal in these assays that was significantly different from the HaloTag® negative control (Fig. 1f), indicating AKAP chains are not in close proximity to each other. This does not preclude AKAP chains being co-localised *in vivo* by interactions with multimeric receptors, e.g. tetrameric TRPV1 (Liao *et al*., 2013).

### Calcineurin binding extends beyond the PxIxIT site

The Calcineurin anchoring site in AKAP5c is at ^337^PIAIIIT^343^, a near-consensus PxIxIT SLiM (Aramburu *et al*., 1998; Garcia-Cozar *et al*., 1998), in which ^338^I takes the position of the consensus proline. In the crystal structure, ^337^P is not engaged, but van der Waals contacts extend ca. five residues C-terminal of the PxIxIT site to ^348^S (Li *et al*., 2012). To investigate the involvement of more distal flanking regions, we titrated Calcineurin into ^15^N-labelled AKAP5c and followed binding by ^15^N HSQC (Fig. 2a). Many peaks showed a progressive decrease in intensity. At one molar equivalent, the reductions were significant, including those expected from residues ^338^IAIIIT^343^ (shown boxed). The changes were purely to the peak intensities; no significant chemical-shift changes were seen, nor did the peaks get broader, indicating that the interaction was in slow exchange on the chemical-shift timescale. In support of this, a few new peaks, presumably from the complex, were discernable close to the noise level but they were extremely broad, indicating slow overall tumbling of the complex (Calcineurin is 80 kDa), rendering most of the residues in the region engaging with Calcineurin invisible. The intensity changes when mapped onto the sequence reveal a binding footprint that extends much further than the crystal structure: approx. 24 residues C-terminal beyond the PIAIIIT site, with further broad dips around residues 390 and 410 (Fig. 2b). Calcineurin binding thus not only extends well beyond the PIAIIIT SLiM but also involves distal regions of the C-terminus of AKAP5c.

**Figure 2:**
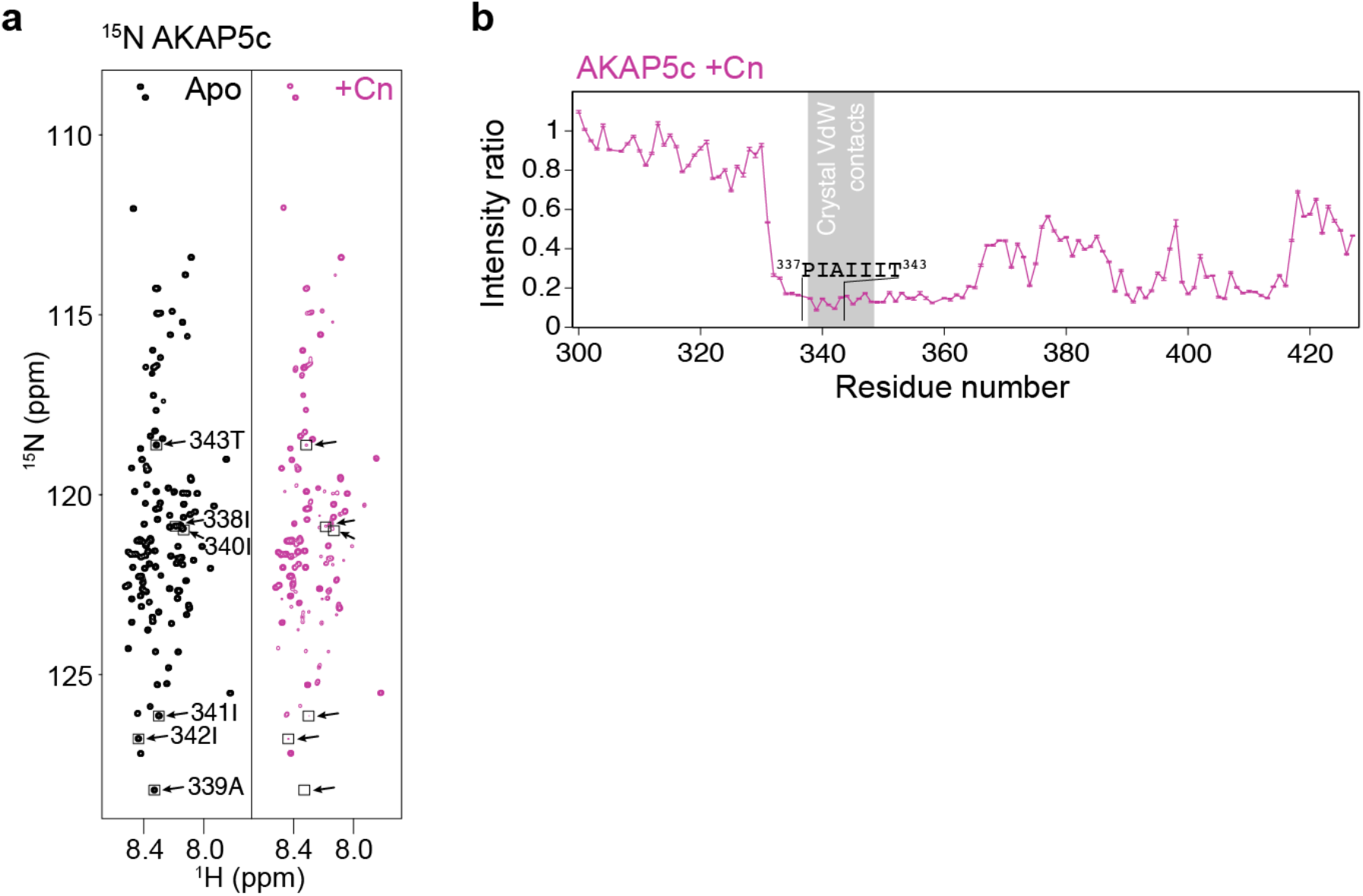
Interactions of AKAP5c with Calcineurin and Calmodulin by ^15^N HSQC NMR. ^15^N-labelled AKAP5c (black) titrated with Calcineurin (Cn; magenta), followed by HSQC. (**a**) HSQC of AKAP5c before and after addition of Cn. Peaks from non-proline residues in ^337^PIAIIIT^343^ are boxed with arrows. (**b**) Intensity changes relative to ^15^N-AKAP5c alone. A molar ratio of 1:1 Cn:AKAP is shown. Grey boxed region: van der Waals contacts in the X-ray crystal structure (3LL8).

### Calcium and Calmodulin do not alter PxIxIT binding

Although the focus of this study is on Calcineurin binding, we wanted to establish whether the known regulators of Calcineurin – calcium and Calmodulin – alter the interaction with PIAIIIT or the new interactions found above. Calcineurin has a regulatory domain that renders it enzymatically dormant in the absence of Ca^2+^ and Calmodulin. Increases in Ca^2+^ concentration lead to partial activation of Calcineurin, through loosening association of the regulatory domain and exposure of the Calmodulin binding region. Further binding of Ca^2+^-Calmodulin results in full activation (Li, Rao and Hogan, 2011; Rumi-Masante *et al*., 2012). In addition to stimulating Calcineurin activity, this exposes a second SLiM-binding region, which interacts with binding partners containing a πϕLxVP motif (π = polar, ϕ = hydrophobic) in a Ca^2+^-Calmodulin dependent manner (Sheftic, Page and Peti, 2016). A titration of the pre-formed AKAP/Calcineurin 1:1 complex with Ca^2+^, followed by Calmodulin, showed that Calcineurin-PIAIIIT binding is independent of both calcium and Calmodulin (Supp. Fig. S3a&b), in line with pull-down experiments from other laboratories (Gold *et al*., 2011; Nygren *et al*., 2017). However, Calmodulin was found to compete with Calcineurin for binding to residues 390-417 of AKAP5c (Supp. Fig. S3c&d), a region ca. 50 residues from the canonical PxIxIT site that coincides with the PKA SLiM (Fig. 1). Interestingly, the sequence is not predicted to be Calmodulin-binding by a database search (Calmodulin Target Database (http://calcium.uhnres.utoronto.ca/ctdb/)). However, it is known to form an amphipathic helix on binding PKA, and contains two clusters of bulky hydrophobic residues separated by a stretch of polar residues, which is a feature of Calmodulin binding (Tidow and Nissen, 2013).

### PxIxIT is not required for binding to non-canonical sites

Given the involvement of PxIxIT-flanking and distal regions of AKAP5c in binding to Calcineurin, we investigated binding to AKAP5c in which the PIAIIIT site had been deleted: AKAP5c_ΔPIAIIIT_. The titration with Calcineurin (Fig. 3a) was carried out as previously. From the overlay of the intensity ratios (Fig. 3b), it is clear that the immediate footprint around the PIAIIIT site (residues 330-350) is lost, as expected given the deletion. However, intensities more distal to the PIAIIIT site still showed significant decreases in intensity, in the same regions and even more pronounced than in WT AKAP5c. Three distinct sections with pronounced dips in intensity were seen, ^361^QFLIS^365, 390^TLLIET^395^ and ^404^IQLSIEQLVN^413^, which we termed secondary sites 1, 2 and 3, respectively.

**Figure 3:**
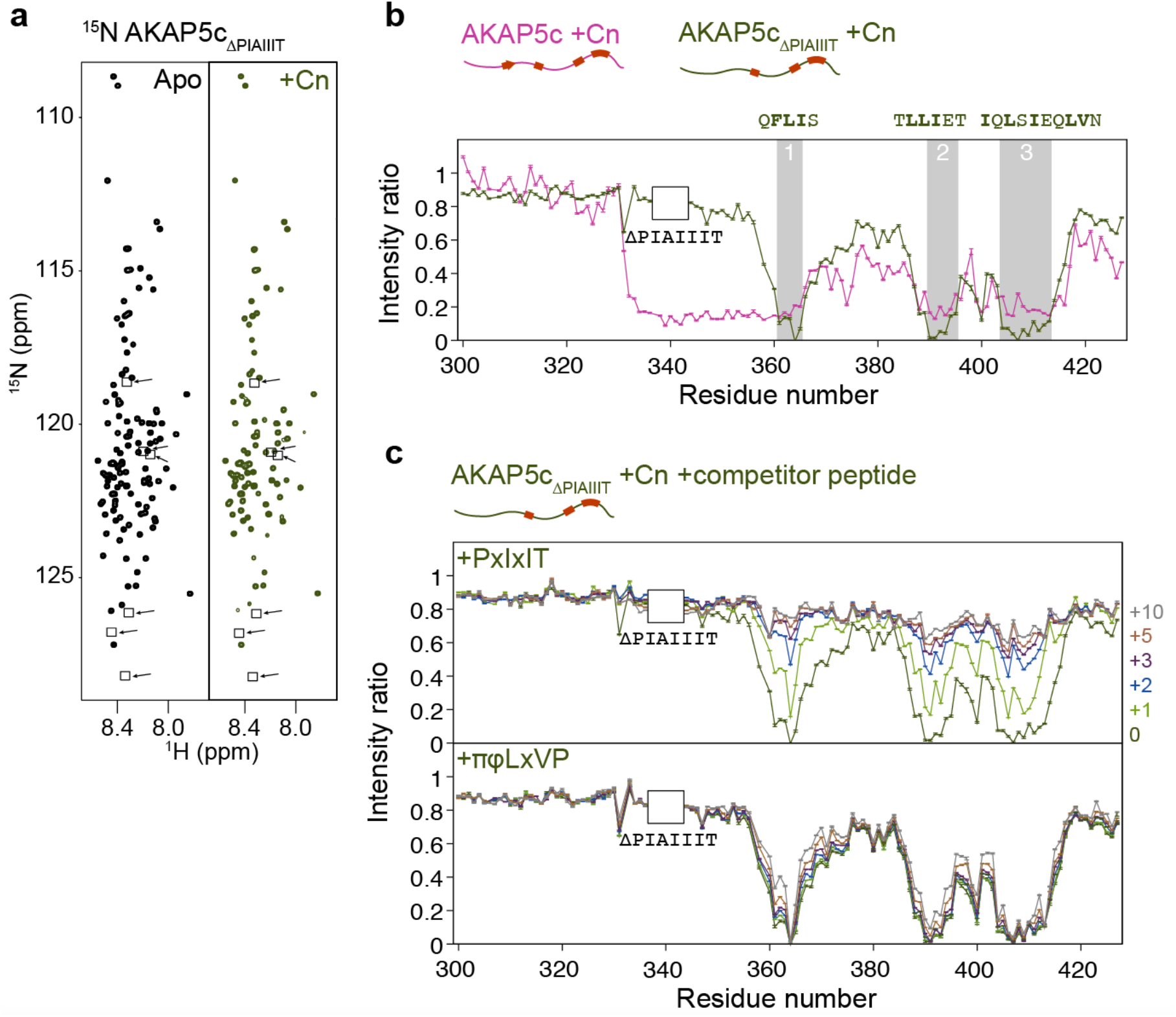
Interactions of AKAP5c_ΔPIAIIIT_ with Calcineurin by ^15^N HSQC NMR. (**a**) ^15^N-labelled AKAP5c_ΔPIAIIIT_ alone (black; positions of deleted ^337^PIAIIIT^343^ residues shown boxed with arrows). AKAP5c_ΔPIAIIIT_ titrated with Calcineurin (Cn) to 1:1 Cn:AKAP (green). (**b**) Intensity changes relative to ^15^N-AKAP5c_ΔPIAIIIT_ alone. Corresponding WT data (magenta) reproduced from Fig. **2b**. Grey boxed regions 1-3 indicate Cn binding sites in AKAP5c_ΔPIAIIIT_, with the corresponding amino acid sequences shown above. (**c**) Titrations of the 1:1 AKAP5c_ΔPIAIIIT_:Cn complex with PxIxIT and πϕLxVP peptides (0 to 10 molar equivalents shown).

Binding to these sites therefore does not depend on the presence of the PIAIIIT site. Moreover, there is some evidence for competition with PIAIIIT since deletion of PIAIIIT appears to enhance the fraction bound at the secondary sites. An important question not resolved by these experiments is whether PIAIIIT and the secondary sites bind to the same or different sites on Calcineurin. In order to address this, the 1:1 ^15^N-AKAP5c_ΔPIAIIIT_/Calcineurin complex was further titrated with a PIAIIIT-containing peptide (E<u>PIAIIIT</u>DTE; Fig. 3c). The peptide proved an effective competitor; ^15^N-AKAP5c_ΔPIAIIIT_ peak intensities were almost fully restored after addition of three molar equivalents of the peptide (Fig. 3c, top). Given the similar hydrophobicity of the secondary sites to the πϕLxVP motif, which binds Calcineurin at a second SLiM-binding region near the heterodimer interface, the experiment was repeated with a πϕLxVP peptide (DD<u>QYLAVP</u>QH (Sheftic, Page and Peti, 2016)). This was ineffective even upon addition of ten molar equivalents (Fig 3c, bottom). The recognition site for secondary sites 1, 2 and 3 is thus the PxIxIT-binding groove in Calcineurin and not the πϕLxVP binding site.

The secondary site sequences each contain three or more bulky hydrophobic residues and include isoleucine, which together are likely to underpin binding to the PxIxIT-binding groove. They are also conserved across PIAIIIT-containing AKAPs (Supp. Fig. 1b). Therefore, a further AKAP mutant was tested: AKAP_ILVF->SA_, in which the PIAIIIT site was present, but all the hydrophobic I, L, V and F residues in the three secondary sites were mutated to S or A to give ^361^QASAS^365, 390^TASAET^395^ and ^404^AQSSAEQSAN^413^. Following reassignment of the chemical shifts, the protein was again titrated with Calcineurin (Fig. 4a). Similar to the WT, large reductions in intensities were observed in and around the PIAIIIT site (Fig. 4b). This was more pronounced in AKAP_ILVF->SA_ indicating higher occupancy of the site by Calcineurin, consistent with removal of competing secondary sites. No dips were seen at secondary sites 2 and 3, (intensities were similar to the unbound AKAP) and site 1 was partially attenuated (overlap with the PxIxIT footprint made the effect less clear). The secondary sites therefore each constitute additional hydrophobic SliMs that despite not being canonical PxIxIT sites, are nevertheless recognised by the PxIxIT-binding pocket on Calcineurin. A small extra region of non-PIAIIIT interaction was apparent in AKAP_ILVF->SA_ in the absence of the secondary sites: ^371^VGVFA^375^, which contains three bulky hydrophobic residues but no isoleucine, and there were also weaker dips corresponding to regions containing just one or two hydrophobic residues: ^378^NG<u>F</u>ED^382, 386^EQ<u>Y</u>ET^390, 398^S<u>LV</u>KN^402^ and ^424^N<u>LL</u>Q^427^ (shown by asterisks in Fig. 4b). We hereon refer to these as ‘mini-SLiMs’ in order to differentiate them from the primary and secondary sites.

**Figure 4:**
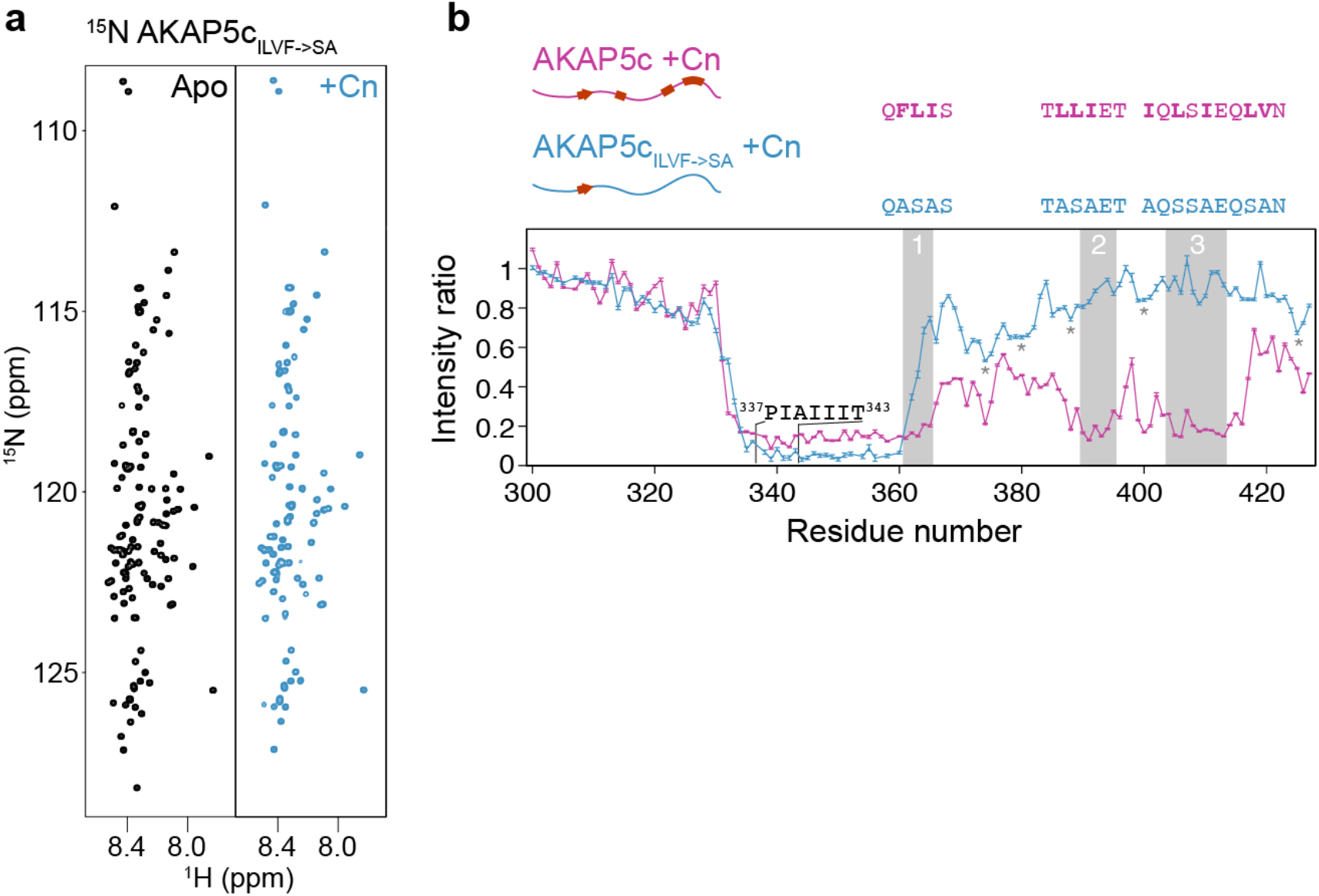
Interactions of AKAP5c _ILVF->SA_ with Calcineurin by ^15^N HSQC NMR. (**a**) ^15^N-labelled AKAP5c_ILVF->SA_ alone (black). AKAP5c_ILVF->SA_ titrated with Calcineurin (Cn) to 1:1 Cn:AKAP (blue). (**b**) Intensity changes relative to ^15^N-AKAP5c_ILVF->SA_ alone. Corresponding WT data (magenta) reproduced from Fig. **2b**. Grey boxed regions 1-3 indicate ‘secondary’ Cn binding sites identified in Fig. **3b**, with the corresponding amino acid sequences shown above. Grey asterisks indicate potential ‘mini-SLiMs’ (see main text).

### Non-canonical sites sequester additional Calcineurin

The detection of three secondary Calcineurin-binding SLiMs led us to speculate about their relative affinity compared to PxIxIT, and whether there is interplay between them. Modified AKAP constructs, Cys-AKAP5c, Cys-AKAP5c_ΔPIAIIIT_ and Cys-AKAP5c_ILVF->SA_ were chemically biotinylated for N-terminal-specific immobilisation on a streptavidin surface so that Calcineurin binding could be assessed by SPR. Calcineurin binding to AKAP5c (Fig. 5a) was clear but complex: the response curves (1) had no clear plateau and (2) displayed evidence of multiple binding sites with both fast and slow kinetics in both the association and dissociation phases. In addition, a fraction of the Calcineurin remained bound at long times. The complexity was reduced somewhat in the curves for AKAP5c_ILVF->SA_ (Fig. 5b), which showed a clear saturation plateau that was absent in the WT, and complete dissociation. For AKAP5c_ΔPIAIIIT_ (Fig. 5c), the curves showed a clear plateau and complete dissociation, but weaker overall binding. Also noteworthy was that the binding curves for AKAP5c_ILVF->SA_ and AKAP5c_ΔPIAIIIT_ did not sum to reproduce the AKAP5c binding curve, i.e. A ≠ B + C, despite their comprising separately the two kinds of binding site present in the WT. This was particularly apparent in the dissociation phase, where the WT showed a sizeable fraction of Calcineurin was essentially permanently bound, while no long-lasting binding of Calcineurin was observed with either mutant.

**Figure 5:**
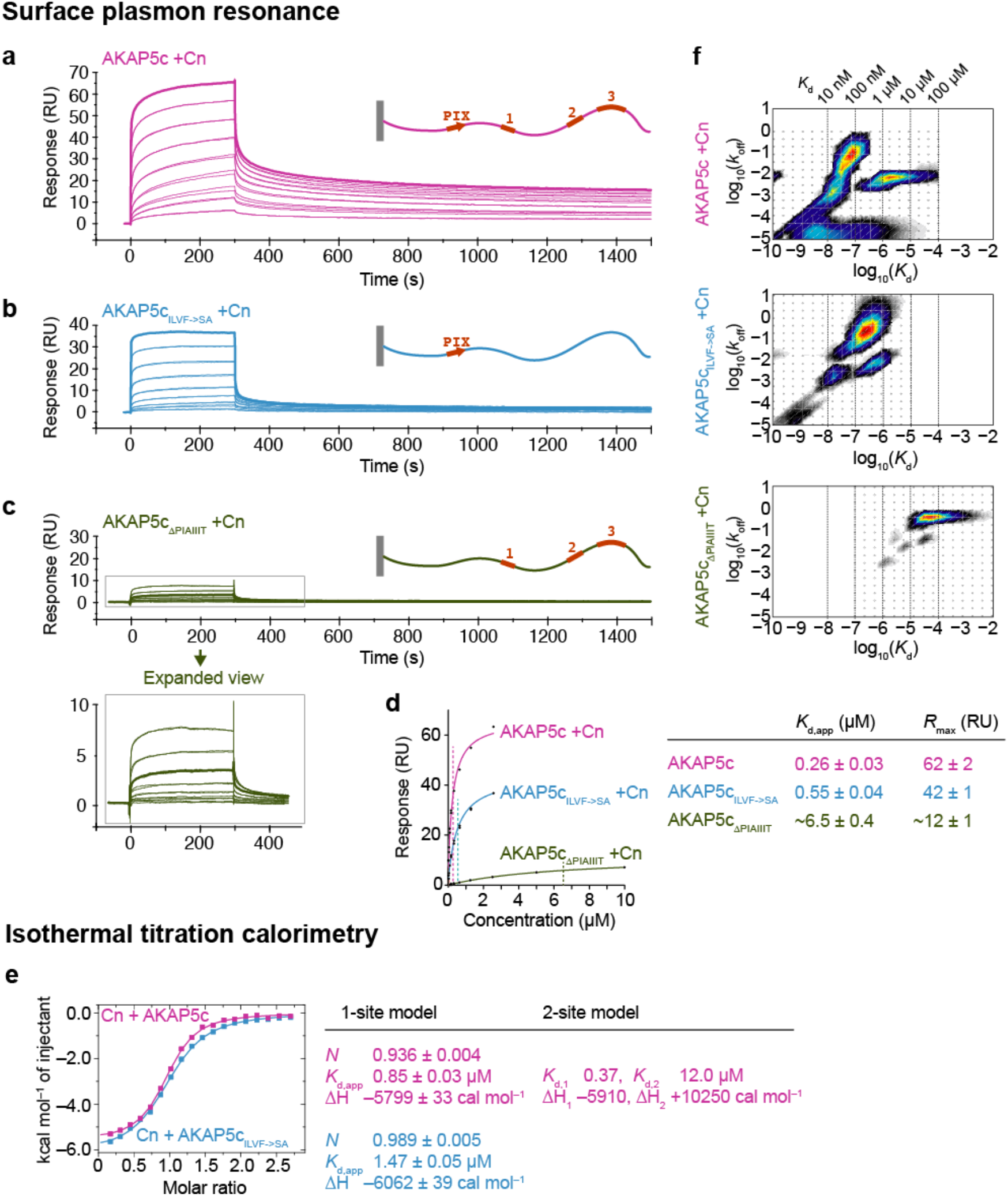
cAKAP/Calcineurin binding by SPR and ITC. Calcineurin injected over a streptavidin surface pre-immobilised with (**a**) AKAP5c, (**b**) AKAP5c_ILVF->SA_ or (**c**) AKAP5c_ΔPIAIIIT_. Calcineurin concentrations were 2-fold dilutions from 2.56 µM (AKAP5c and AKAP5c_ILVF->SA_) or 10 µM (AKAP5c_ΔPIAIIIT_), in duplicate. (**d**) Fits of ‘steady-state’ response vs Calcineurin concentration used to obtain estimates of *K*_d,app_ and *R*_max_. (**e**) ITC of AKAPs injected into Calcineurin gives sigmoidal exotherms that can be fit to a 1-site model. Following the observation of lower stoichiometry for AKAP5c (*N* = 0.936), the data were also fit to a 2-site model with fixed *N* = 1 stoichiometries. **(f)** SPR surface site complexity modelled in Evilfit (Schuck and Zhao, 2010): distribution plots of *k*_off_ vs *K*_d_ obtained by fitting the data from **a**-**c** using a regularization and Bayesian approach (see Supp. Fig. S4 for further details).

An estimate of the *K*_d_ could be obtained from fitting the signal (RU) at the end of the association period *vs* Calcineurin concentration (Fig. 5d). The values obtained can only be estimates for AKAP5c and AKAP5c_ΔPIAIIIT_ due to the lack of saturation (‘steady-state’) of the former and the lack of high-concentration data for the latter. Nevertheless, the trend in apparent *K*_d_ is clear: AKAP5c binds Calcineurin about twice as tightly compared to AKAP5c_ILVF->SA_, (*K*_d_ ∼0.3 vs 0.6 µM), and both PIAIIIT-containing AKAP constructs bind Calcineurin with at least ten-fold higher affinity than AKAP5c_ΔPIAIIIT_, which has a *K*_d_ of ∼ 10 µM. Estimation of the total surface capacity for Calcineurin (*R*_max_) is also instructive as it gives information on the stoichiometry of binding under saturating conditions. From extrapolation of the binding curves, it appears that AKAP5c can accommodate significantly more Calcineurin than AKAP5c_ILVF->SA_ (62 vs 42 RU; Fig. 5d). The secondary sites appear to account for at least some of this ‘extra’ capacity, since *R*_max_ for AKAP5c_ΔPIAIIIT_ is (very approximately) 12 RU.

Binding was also investigated by isothermal titration calorimetry (ITC). Titration of AKAP5c and AKAP5c_ILVF->SA_ into Calcineurin gave clear sigmoidal binding curves (Fig. 5e). Low heats prevented detection of binding to AKAP5c_ΔPIAIIIT_ (not shown), presumably due to insufficient binding at the attainable concentrations of free Calcineurin or small ΔH, or both. The data for AKAP5c and AKAP5c_ILVF->SA_ both fit well to a 1-site model, but with some differences: the stoichiometry (*N*) was consistently slightly lower for AKAP5c over many repeats (0.94 vs 0.99 AKAP5c:Cn), while the *K*_d_ was approx. two-fold tighter (0.85 vs 1.5 µM). Small differences were also seen in ΔH, AKAP5c_ILVF->SA_ being slightly more exothermic. Given that the < 1 stoichiometry was likely to be due to the presence of competing secondary binding sites on cAKAP5 (Fig. 3), the data were also fit to a 2-site model with fixed *N* = 1 stoichiometries (Zhao, Piszczek and Schuck, 2015), which yielded a fit of the same quality but two distinct *K*_d_s (0.37 and 12 µM). The presence of two or more sites is not inconsistent with the *N* = 0.94 obtained from the 1-site fit, since the second, 30-fold weaker binding site would not be well populated at the concentrations possible for ITC (18 µM), especially when competing for Calcineurin with a much tighter site.

The multivalency of AKAP5c accounts for the deviation from ideal pseudo-first order binding kinetics seen by SPR (Fig. 5a). Such systems require a different analysis strategy if they are to be properly characterised. A suitable alternative approach has been developed by Schuck and colleagues (‘Evilfit’ (Schuck and Zhao, 2010)), which uses modern regularization methods combined with a Bayesian approach to add features that are essential to explain the observed data to the single species model. Such features could be populations of lower-affinity or poorly reversible sites. With this approach, it was possible to obtain excellent fits to the raw SPR data (Supp. Fig. S4), and to extract *k*_off_ vs *K*_d_ distribution plots (Fig. 5f). The complexity seen for Calcineurin binding to AKAP5c was deconvolved into 3 or 4 classes of surface sites: one or possibly two very similar sites with *K*_d_ 10-100 nM, another with *K*_d_ 1-10 µM, and a final class with a very slow *k*_off_ that presumably accounts for the lack of full dissociation observed even at long times (i.e. poor reversibility). The number of surface site classes was reduced in AKAP5c_ILVF->SA_ to one main site with *K*_d_ 100 nM-1 µM. The distribution was simpler again for AKAP5c_ΔPIAIIIT_ which displayed a single class of sites with *K*_d_ 10-100 µM (the true value is likely to lie at the lower end of this range given the values obtained from the equilibrium fit (∼10 µM; Fig. 5d), by ITC (∼12 µM; Fig. 5e) and the fact that a high fraction bound was seen by NMR at a concentration of 100 µM (Fig. 3b)). These results support a picture in which AKAP5c acts as a multivalent surface capable of binding one or more molecules of Calcineurin at different sites.

The ‘long-lived’ (slow *k*_off_) bound population is only significant when both sites are present (Fig. 5a&f), and is therefore dependent on a multivalent AKAP. However, it is possible that Calcineurin contributes additional multivalency. We observed a weak propensity for formation of heterotetramers by AUC (8% at 7.5 µM; Supp. Fig. S5a), indicating a dimer/tetramer *K*_d_ in the 100 µM range, well above the concentrations used for SPR (10 nM - 2.56 µM). However, this was even lower in the complex (6%; Supp. Fig. S5b), indicating that tetramerisation was either slightly reduced by AKAP binding, or replaced with a small amount of purely AKAP-mediated 2:1 Cn:AKAP complex, or both (unfortunately it is not possible to distinguish the two due to the small percentage and compensatory effects of higher friction and higher mass). This was corroborated by our observations that the aggregation tendencies of Calcineurin, which can be severe at high concentrations or lower pHs, were substantially alleviated when in complex with a PxIxIT or secondary motif-containing substrate, which suggests that the aggregation tendency is mediated by an unmasked PxIxIT-binding groove. At the high concentrations used for NMR (75 µM) the higher *s*-value species increased to 18% (Supp. Fig. S5c). The impact of heterotetramers was further explored by repeating the NMR and SPR experiments in the presence of a large excess of DTT, which disfavours disulphide-mediated oligomerisation (Calcineurin contains 14 cysteines, many of which are solvent exposed). Both the AKAP5c/Cn NMR intensity changes and the long-lived population of Calcineurin seen by SPR were preserved (Supp. Fig. S6). We conclude that the slow observed *k*_off_ phenomenon seen by SPR hinges on multivalency of AKAP, rather than Calcineurin, and is therefore reminiscent of the re-capture behaviour seen for antigens binding to bivalent antibodies, which also show anomalously slow dissociation, or ligands rebinding to receptors at cell surfaces (Lagerholm and Thompson, 1998).

## Discussion

From our experiments, it is clear that the Calcineurin binding sites in AKAP5c are far more extensive than PIAIIIT, including additional SLiMs and ‘mini-SLiMs’ in the C-terminal region that interact with µM-mM affinity: non-consensus SLiMs containing as few as three bulky hydrophobic residues appear able to bind the PxIxIT-groove with low-to-mid µM affinity (AKAP5c_ΔPIAIIIT_; Fig. 5d&f), and mini-SLiMs containing only one or two hydrophobic residues also produce detectable binding by ^15^N-HSQC (Fig. 4b). These additional sites increase both the apparent affinity, and potentially the binding capacity of the scaffold. The apparent affinity is probably raised through both an increase in the number of productive encounters given the high density of SLiMs, and through rebinding mechanisms that reduce the diffusion of Calcineurin away from the scaffold since the molecule is likely to be intercepted by another binding site rather than diffusing away. This latter point is borne out directly in our experiments by the lack of complete dissociation of Calcineurin on AKAP5c seen by SPR (Fig. 5a). The ‘labile but localised’ pool of rapidly dissociating, diffusing and rebinding Calcineurin can presumably also be captured by downstream effectors such as NFAT despite their mid µM affinities (Fig. 6).

**Figure 6:**
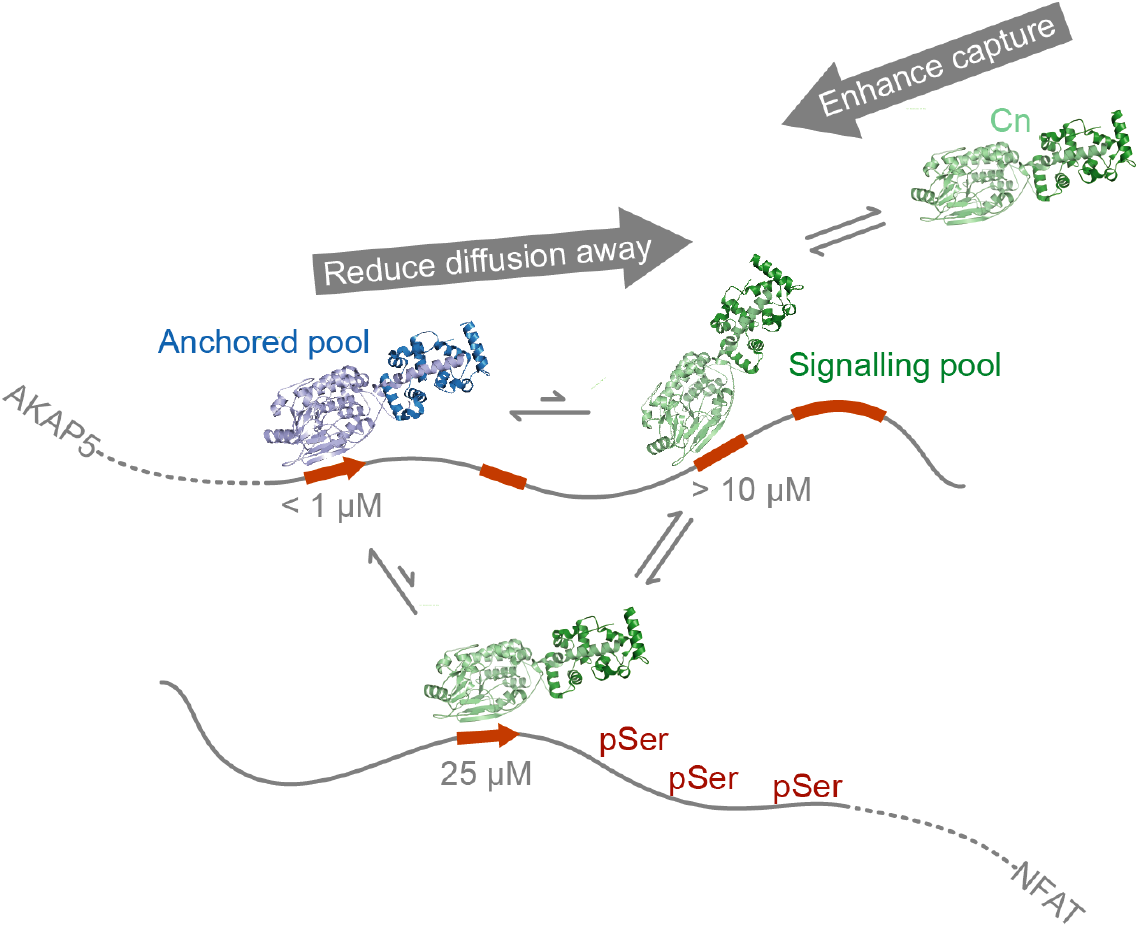
Non-canonical SLiMs could facilitate both anchoring and formation of a labile Calcineurin pool that enables efficient transfer to effectors. AKAP5c captures and retains Calcineurin using its consensus PIAIIIT (red arrow) and secondary sites (red rectangles), building anchored and signalling pools of bound Calcineurin (blue and green, respectively). Calcineurin binds with lower-affinity to the secondary sites, enabling effective competition by weaker PxIxIT motifs on downstream effectors, e.g. the PRIEIT SLiM in NFAT, and optimal signalling.

These new insights are made possible by the combination of NMR experiments and a non-traditional approach to SPR data fitting that is validated by strategic mutagenesis. The experimental study of molecular recognition by disordered proteins can be very challenging due to the rapid increase in complexity inherent in multivalent binding. Here we demonstrate that weak binding by SLiM flanking regions can give rise to the non-integer stoichiometries often seen experimentally by ITC, and SPR binding curves with a high degree of heterogeneity, both of which can be difficult to rationalise and are often dismissed as artefacts due to incorrect concentrations/activities in the case of ITC, or orientational/crowding effects on the SPR surface. These phenomena – even if relevant to the biology as here – can limit the study of multivalent complexes to their global behaviour only, with the loss of mechanistic information. An alternative is dissection into simpler pairwise interactions, treating the intervening sequences as inert (which may incur the loss of important information on competition, cooperativity and multiple binding modes). SPR data on multivalent systems are often high quality in both signal-to-noise and reproducibility, while being deemed “unfittable” due to the ill-posed problem of decomposing a distribution of exponentials. However we find that the combination of regularization and Bayesian approaches (Schuck and Zhao, 2010), designed for the deconvolution of mass transport and surface heterogeneity, is also very effective at decomposing multivalent interactions. The resulting distributions may be subsequently validated by competition and mutagenesis, as we have done here, alongside mapping by NMR, which is both residue-specific and sensitive to a wide range of interaction affinities (Zuiderweg, 2002).

We find that the PxIxIT-binding groove recognises hydrophobic residues, particularly isoleucine, and also appears to mediate further oligomerisation of Calcineurin when vacant. However, further evidence for weak, non-canonical engagement of the groove with hydrophobic residues is seen in the existing PxIxIT peptide/Calcineurin X-ray crystal structures, in which the reverse face of the PxIxIT site provides an additional potential encounter point (Supp. Fig. S7a). In the structures, one with PVIVIT (Li *et al*., 2007) and one with the AKAP sequence PIAIIIT (Li *et al*., 2012), the stoichiometry in the asymmetric unit is two Calcineurin heterodimers to one peptide, with the second heterodimer binding to the opposite side of the PxIxIT site, out of register by one residue towards the C-terminus of AKAP5. The peptide thus forms an axis of pseudosymmetry. In both cases, the first heterodimer that engages the consensus elements of the PxIxIT sequence has a larger buried surface area and a greater number of hydrogen bonds. The opposite side displays VxV or AxI, to which a second heterodimer is bound in the crystal, using the same hydrophobic PxIxIT-binding groove, but with significantly lower affinity, since no evidence for a 2:1 stoichiometry was seen in solution by size exclusion chromatography or SEC-MALS (Li *et al*., 2007, 2012). However, the presence of hydrophobic residues in the two ‘x’ positions may serve as an additional capture surface on PxIxIT, even if unstable. Their presence may (partially) account for the fact that in our experiments, mutation of the three distal secondary sites in AKAP5c_ILVF->SA_ did not restore ‘pure’ pseudo first-order binding characteristic of simple bimolecular collisions (Fig. 5b & f). Considered as a group, PxIxIT affinities span a wide range, and the fine-tuning of PxIxIT affinities by altering the ‘x’ positions presumably facilitates evolution into either anchoring or signalling roles. It is interesting to note that during selection for competitive peptide inhibitors of PxIxIT binding there was a clear enrichment for hydrophobic residues (V, I and L) in the two x positions (Aramburu *et al*., 1999), while many of the low affinity substrates favour charged or polar amino acids in these positions that would be incompatible with the non-polar groove on Calcineurin e.g. PRIEIT in NFATc1 (25 µM; Supp. Fig. S7b).

Our findings also demonstrate that SLiM recognition in AKAP5c is non-exclusive: both Calcineurin and Calmodulin recognise features of the canonical PKA SLiM and therefore all three are in competition for the same site (Fig. 2 & 3). Interactions of one SLiM with many partners have been documented (Hsu *et al*., 2013), and due to the substitution-tolerant nature of SLiMs coupled with the conformational plasticity of IDPs, may be a common feature of disordered scaffolds. In this example, PKA has a higher affinity for AKAP5c than Calcineurin (and presumably Calmodulin). However, the *in vivo* concentration of cellular PKA is sub-µM (Beavo, Bechtel and Krebs, 1974) whereas concentrations of Calcineurin are generally much higher (e.g. 1% of the total protein concentration in brain (Yakel, 1997)), and further concentrated at the membrane or cytoskeletal elements (Shibasaki, Hallin and Uchino, 2002). A slowly diffusing pool of highly concentrated Calcineurin (or Calmodulin), fed by an intrinsically high cellular presence and maintained by other local SLiMs, will have a high effective concentration and therefore be competitive. Moreover, due to the high abundance of AKAPs, there is likely to be an excess of anchoring sites compared to the kinase (Omar and Scott, 2020).

Parallels can be drawn between a ‘labile but localised’ pool of Calcineurin that supplies the canonical anchoring site and the ways transcription factors (TFs) bind DNA. Here, the capture is different in that it involves two proteins, and a patchy hydrophobic surface rather than the smooth negatively charged phosphate backbone of DNA. TFs recognize their binding site with high affinity, but bind with low affinity to non-canonical DNA, which reduces the dimensionality of the binding-site search *via* mechanisms such as sliding, jumping and monkey bar strand transfer (Gowers and Halford, 2003; Vuzman, Azia and Levy, 2010). In the case of AKAP5c, which unlike DNA is inherently flexible, an additional mechanism could be invasion and competitive displacement of secondary SLiMs by PxIxIT, enhanced by its high effective molarity. The recognition of IDPs containing multiple SLiMs for a partner has been termed ‘allovalency’ (Klein, Pawson and Tyers, 2003; Levchenko, 2003). In this example, the SLiMs have very different affinities for the partner and yet the mM-affinity partial/mini-SLiMs may be essential to conserve the hydrophobic ‘capacitance’ of the region responsible for Calcineurin capture.

In conclusion, here we uncover several Calcineurin interaction motifs spanning a wide range of affinities in a short region of a scaffold protein. At least three hydrophobic patches bind the Calcineurin PxIxIT groove with µM affinity, which is similar to Calcineurin’s affinity for many of its PxIxIT-containing substrates. The patches in AKAP5c are conserved across relevant AKAPs (Supp. Fig. S1b) and do indeed appear to bestow a biologically useful function: to enhance capture and anchoring, while maintaining a more mobile pool of Calcineurin than would be present if the anchoring protein contained one high affinity SLiM, thus acting together to maintain a stable but responsive signalosome.

## Materials and Methods

### Constructs

pcDNA™3.1 V5-His TOPO containing full length human AKAP5 with V5 and His tags was obtained. AKAP5c_ILVF->SA_ was generated by gene synthesis (GeneArt; ThermoFisher). pOP5T was created from pOP5TT (obtained from Marko Hyvönen, Department of Biochemistry, University of Cambridge), by deletion of thioredoxin leaving a TEV-removable N-terminal octa-His tag. pET15b CnA CnB, which contains human PPP3CA and PPP3R1 (Mondragon *et al*., 1997) was obtained from Addgene (11787). Plasmid pQE30 CaM containing rat Calmodulin (protein sequence 100% identical to human) was obtained from Dermot Cooper, Department of Pharmacology, University of Cambridge. NanoBiT vectors were obtained from Promega.

Cloning was carried out using standard molecular biology techniques. The coding sequence for AKAP5c was PCR amplified from pCDNA3.1 hAKAP and inserted between the BamHI and HindIII sites of pHAT3 (Peränen *et al*., 1996) using restriction cloning. All other cloning was carried out using a modified SLIC protocol based on (Jeong *et al*., 2012). The appropriate plasmid and insert sequences (see above) with 15 bp complementary overhangs, or 12 bp for deletion of the PIAIIIT site were amplified using Q5 polymerase (NEB). For unique Cys mutants, a Cys codon was added to the forward primer immediately after the GS in the Thrombin or TEV site.

Final constructs were:

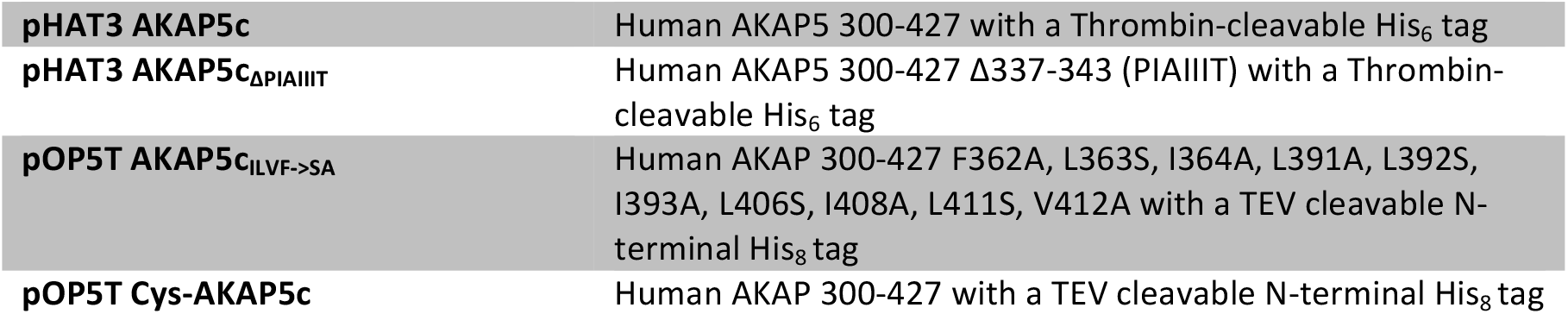
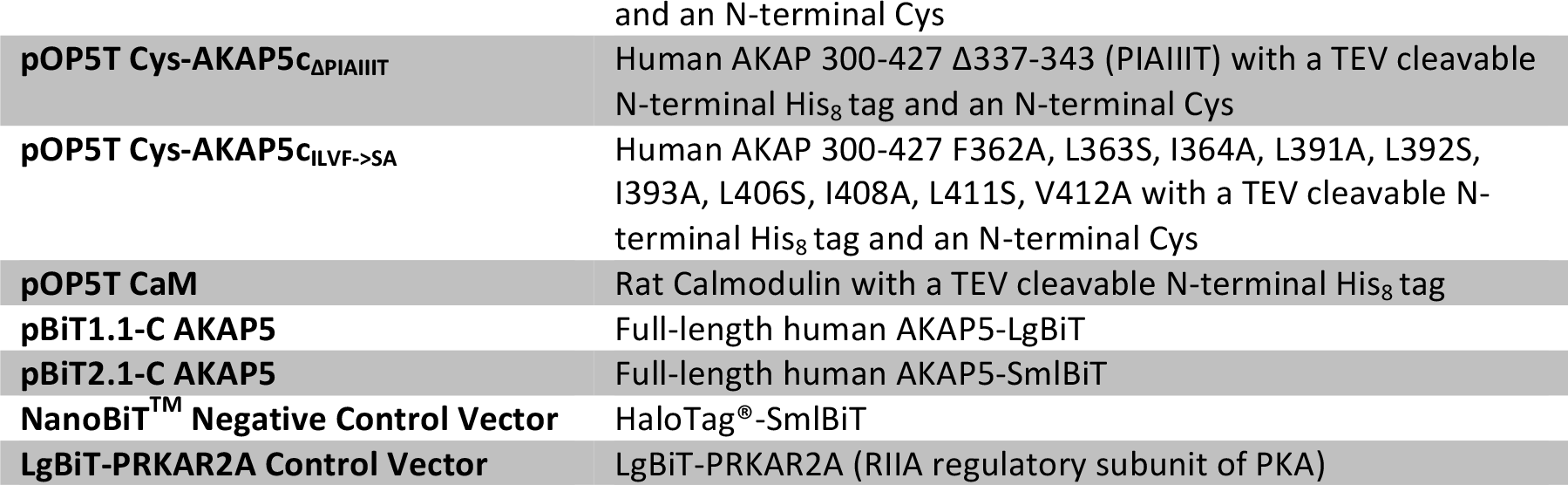

Protein sequences are presented in full in *Supplementary Information*.

### Protein expression and purification

All proteins were expressed in *E*.*coli* BL21(DE3) or BL21 Star (DE3) cells transformed with the appropriate plasmid. For unlabelled proteins 2 L baffled flasks containing 500 ml of AIM LB or AIM 2YT autoinduction medium (Formedium) supplemented with 50 µg/ml carbenicillin (and 1 mM CaCl_2_ for Calmodulin) were inoculated with 1% starter culture in LB medium and grown overnight at 30 °C with shaking at 170 rpm. Labelled proteins were expressed similarly, but in M9 minimal medium containing (as appropriate) 0.4 g/L ^15^N ammonium chloride, 2 g/L uniformly labelled ^13^C glucose as the sole nitrogen and carbon sources and supplemented with 5% Celtone complete medium with the appropriate isotopic labelling. Cells were grown to an OD_600_ of 0.7 before induction with 0.5 mM IPTG, and grown overnight at 23 °C.

AKAP and Calmodulin constructs expressed from either pHAT3 or pOP5T were purified as follows: cells were resuspended in IMAC buffer A (20mM Tris pH 8.0, 0.5 M NaCl, 10 mM imidazole) with SIGMA*FAST* EDTA free protease inhibitor cocktail. Cells were lysed using an Emulsiflex and clarified lysate was heated at 70 °C for 20 min. Precipitated protein was removed by centrifugation and the supernatant further purified by Ni-NTA affinity chromatography. Bound proteins were eluted with IMAC buffer B (A containing 400 mM imidazole) and dialysed overnight into 20 mM Tris pH 8.0 using 7 kDa cut-off membrane, the His tag was removed with thrombin (pHAT3) or TEV (pOP5T), uncut protein removed by passing over Ni-NTA resin, and proteins further purified using a 6 ml Resource Q column (Cytiva) eluted with a linear NaCl gradient 0-400 mM. DTT (2 mM) was present in all steps after the initial Ni-NTA purification of proteins containing a unique cysteine. Proteins were concentrated and buffer exchanged into the appropriate buffer before storage at –80 °C.

Calcineurin was purified similarly, except that rather than heating, it was precipitated from clarified lysate by addition of ammonium sulphate to 45% saturation on ice before being resuspended in IMAC buffer A, and the His tag on the A subunit was not removed. After elution from the Resource Q column, Calcineurin was concentrated to ∼ 5-10 mg/ml and further purified by size exclusion chromatography using an S200 16/600 column (Cytiva) in 20 mM Tris/HCl pH 7.4, 150 mM NaCl, 2 mM DTT and stored at –80 °C.

### Peptides

Peptides (EPIAIIITDTE and DDQYLAVPQH) were purchased from Biomatik and dialysed extensively against the appropriate buffer before use.

### Biotin labelling

Proteins were labelled using EZ-Link™ Maleimide-PEG2-Biotin (ThermoFisher) in 10 mM PIPES pH 6.5, 150 mM NaCl according to the manufacturer’s instructions.

### Concentration measurements

Protein and peptide concentrations were determined spectophotometrically using a NanoDrop One^C^ instrument (ThermoFisher) at 205 nm for the AKAP constructs (due to their low ε_280_) and 280 nm for Calcineurin and Calmodulin. Extinction coefficients were calculated using https://spin.niddk.nih.gov/clore/Software/A205.html based on Anthis & Clore, 2013 (Anthis and Clore, 2013).

### Nuclear Magnetic Resonance

NMR measurements were made on ^15^N- or ^13^C,^15^N-labelled proteins (50-100 µM) in 10 mM PIPES pH 6.5, 150 mM NaCl (NMR buffer) and 10-15% ^2^H_2_O. Experiments were recorded at 25 °C on Bruker DRX600 or 800 spectrometers. Assignments were obtained using a conventional triple-resonance approach (HNCA, HN(CO)CA, HNCO, HNCACB, HN(CO)CACB) alongside TOCSY-^15^N-HSQC, NOESY-^15^N-HSQC and HNN/HN(C)N experiments (Panchal, Bhavesh and Hosur, 2001; Cavanagh *et al*., 2006). Data were processed using AZARA (v.2.7, © 1993-2022; Wayne Boucher and Department of Biochemistry, University of Cambridge). Triple resonance experiments were recorded with 25% nonuniform sampling, using Poisson-gap sampling (Hyberts, Takeuchi and Wagner, 2010), and reconstructed using the Cambridge CS package and the CS-IHT algorithm (Bostock, Holland and Nietlispach, 2012). Assignments were made using CcpNmr Analysis v. 2.4 (Vranken *et al*., 2005). Chemical shifts were referenced to 2,2-dimethyl-2-silapentane-5-sulfonic acid (DSS). Heteronuclear NOE values were obtained at 600 MHz with either 4 s of ^1^H saturation using a 120° pulse train or a 4 s delay prior to the first ^15^N pulse (Farrow *et al*., 1994). Complexes with Calcineurin were first prepared at the defined stoichiometries in 10 mM Tris/HCl pH 7.4, 150 mM NaCl, 2 mM DTT and then dialysed into NMR buffer. Peptides were added from concentrated stocks to minimise dilution effects. Intensity ratios were calculated using fitted peak heights in Analysis. Chemical-shift differences were calculated using Δδ = [(Δδ^H^)^2^ + (0.15 × Δδ^N^)^2^]^1/2^ (Zuiderweg, 2002).

### Far-UV Circular Dichroism Spectroscopy

Experiments were performed at 0.1 mg/ml over a 185-260 nm range at 25 °C in 10 mM PIPES pH 6.5 and 150 mM NaF and in 1 mm path-length cuvettes. Spectra were acquired using an AVIV 410 spectrometer in 1 nm wavelength steps, averaged over three accumulations and baseline-corrected using buffer before smoothing, using the manufacturer’s software. Millidegree units were converted to mean residue ellipticity (MRE) with units deg cm^2^ dmol^−1^ res^−1^ using MRE = millideg. / {(no. residues – 1) × c × *l* × 10}, where c = molar concentration and *l* = path length in cm.

### Analytical Ultracentrifugation

Sedimentation velocity was measured in an Optima XL-I centrifuge (Beckman Coulter) by interference using standard 12 mm double-sector Epon centrepieces with sapphire windows in an An60 Ti four-hole rotor at 20 °C. Multi-component sedimentation coefficient *c*(*s*) distributions were obtained from by direct boundary modelling of the Lamm equation using Sedfit v.15 (Schuck, 2000). The density and viscosity of the buffers and the partial specific volumes of the proteins were calculated using Sednterp (Laue *et al*., 1992). For AKAP5c, 100 scans were collected over 22 h on 400 µl of AKAP5c at 70 µM or buffer (10 mM sodium phosphate pH 6.5, 150 mM NaCl) at 40 krpm. For Calcineurin and Calcineurin/AKAP5c complex (Supp. Fig. S5 a&b), 64 scans were collected over 5 h on 400 µl of Calcineurin or Calcineurin/AKAP5c (Cn 7.5 µM, AKAP 5 µM) or buffer (10 mM Tris/HCl pH 7.4, 150 mM NaCl, 5 mM DTT) at 50 krpm. Calcineurin/AKAP5c complex at high concentration (Supp. Fig. S5c) was prepared as for the NMR samples but without ^2^H_2_O. 64 scans were collected over 3 h on 130 µl of AKAP5c/Calcineurin at 75 µM or buffer (10 mM PIPES pH 6.5, 150 mM NaCl) at 50 krpm.

### Cell culture

HEK293T cells were cultured in 10 cm tissue culture treated dishes grown at 37 °C in a 5% CO_2_ atmosphere in HEPES buffered DMEM/F12 1:1 (Merck) supplemented with 10% fetal bovine serum and 2 mM L-glutamine. Cells were maintained in logarithmic growth phase and passaged upon reaching ∼80% confluence (every 2-3 days).

### Split-Luciferase Reporter Assays (NanoBiT)

Split luciferase reporter assays were carried out using the Promega NanoBiT system. Briefly, cells were diluted in in HEPES buffered DMEM/F12 1:1 (Merck) supplemented with 10% fetal bovine serum and 2 mM L-glutamine to 50,000 cells/ml and plated at a density of 5000 cells per well (100 µl) in 96-well white clear-bottom tissue culture treated plates (Corning 3610). Cells were grown for 24 h before transfection. Cells were transiently transfected with 50 ng each of the appropriate plasmids using FuGENE HD (Promega) at a FuGENE DNA ratio of 3:1. After transfection cells were grown for a further 24 h before assay.

Plates were left to equilibrate at room temperature for 20 minutes before addition of NanoGlo live cell substrate (Promega) according to the manufacturer’s instructions. After mixing at 800 rpm for 10 s, luminescence was assayed in a PHERAstar microplate reader (BMG LABTECH) at 25 °C using bottom optics and an averaging time of 10 s.

### Surface Plasmon Resonance

Experiments were performed on a Biacore T200 instrument (GE) at 25 °C. Biotin-Cys-AKAP5c, Biotin-Cys-AKAP5c_ILVF->SA_ and Biotin-Cys-AKAP5c_ΔPIAIIIT_ were sparsely immobilised on separate channels of an SA sensor chip (Cytiva). In order to optimise the analyte response signal-to-noise ratio, 12 RU were immobilised for AKAP5c and AKAP5c_ILVF->SA_, and 55 RU for AKAP5c_ΔPIAIIIT_; analyte responses for AKAP5c_ΔPIAIIIT_ were subsequently re-scaled by a factor of 12/55 for comparative purposes. Binding to AKAP5c and AKAP5c_ILVF->SA_ was determined in 10 mM PIPES pH 6.5, 150 mM NaCl, 0.05% v/v Tween20, at ten Calcineurin concentrations (0, 10, 20, 40, 80, 160, 320, 640, 1280 and 2560 nM). Binding to AKAP5c_ΔPIAIIIT_ was determined at nine Calcineurin concentrations (0, 78, 156, 313, 625, 1250, 2500, 5000 and 10000 nM). All concentrations were run in duplicate. The contact time, dissociation time and flow rate were 300 s, 1200 s and 30 µL min^−1^, respectively. The chip surface was regenerated with 50 mM NaOH, 1 M NaCl for 1 or 2 × 60 s. Data were processed using the manufacturer’s BIAevaluation software and further analysis was carried out in EVILFIT (Schuck and Zhao, 2010).

### Isothermal Titration Calorimetry

Experiments were performed on an ITC200 (GE/Malvern) at 25 °C. 250 µM AKAP5c or AKAP5c_ILVF->SA_ in the syringe was injected into 18.32 µM Calcineurin in the cell in 18 × 2 µL steps (after 1 × 0.4 µL), with a stirring speed of 600 rpm and an injection spacing of 120 s. All proteins were dialysed into 10 mM Tris/HCl pH 7.4, 150 mM NaCl, 5 mM DTT prior to use. Data were integrated and analysed in Origin (1-site model), or NITPIC and SEDPHAT (2-site model: ‘A+B+B; two non-symmetric sites’) (Keller *et al*., 2012; Brautigam *et al*., 2016). The fitted parameters following analysis with the 1-site model are the average of two repeats, while those from the 2-site model are the result of a global fit.

## Data availability

Chemical-shift assignments for AKAP5c (see also Supp. Fig. S2) have been deposited in the BMRB with accession number XXXX.

## Acknowledgements

We thank Peter Schuck, Marko Hyvönen and Anna Git for helpful discussions, the NMR and Biophysics Facilities in the Department of Biochemistry, University of Cambridge for access to instrumentation, and Dermot Cooper for the Calmodulin plasmid. This work was supported by the Biotechnology and Biological Sciences Research Council (BB/N022181/1).

