## Supplementary Information for "Hidden multivalency in phosphatase recruitment by a disordered AKAP scaffold"

### APPENDIX

#### Protein sequences

AKAP5C
GSVAEETKPKDTELSQESDFKENGITEEKSKSEESKRMEPIAIIITDTEISEFDVTKSKNVP
KQFLISAENEQVGVFANDNGFEDRTSEQYETLLIETASSLVKNAIQLSIEQLVNEMASDDNK
INNLLQ
$\varepsilon_{205}$= 420860 $M^{-1}$ $cm^{-1}$

AKAP5C$_{\Delta PIAIIIT}$
GSVAEETKPKDTELSQESDFKENGITEEKSKSEESKRMEDTEISEFDVTKSKNVPKQFLISA
ENEQVGVFANDNGFEDRTSEQYETLLIETASSLVKNAIQLSIEQLVNEMASDDNKINNLLQ
$\varepsilon_{205}$= 401400 $M^{-1}$ $cm^{-1}$

AKAP5C$_{ILVF\text{->}SA}$
GSVAEETKPKDTELSQESDFKENGITEEKSKSEESKRMEPIAIIITDTEISEFDVTKSKNVP
KQASASAENEQVGVFANDNGFEDRTSEQYETASAETASSLVKNAAQSSAEQSANEMASDDNK
INNLLQ
$\varepsilon_{205}$= 412260 $M^{-1}$ $cm^{-1}$

Cys-AKAP5C
GSCVAEETKPKDTELSQESDFKENGITEEKSKSEESKRMEPIAIIITDTEISEFDVTKSKNV
PKQFLISAENEQVGVFANDNGFEDRTSEQYETLLIETASSLVKNAIQLSIEQLVNEMASDDN
KINNLLQ
$\varepsilon_{205}$= 424330 $M^{-1}$ $cm^{-1}$

Cys-AKAP5C$_{\Delta PIAIIIT}$
GSVAEETKPKDTELSQESDFKENGITEEKSKSEESKRMEDTEISEFDVTKSKNVPKQFLISA
ENEQVGVFANDNGFEDRTSEQYETLLIETASSLVKNAIQLSIEQLVNEMASDDNKINNLLQ
$\varepsilon_{205}$= 404870 $M^{-1}$ $cm^{-1}$

Cys-AKAP5C$_{ILVF\text{->}SA}$
GSVAEETKPKDTELSQESDFKENGITEEKSKSEESKRMEPIAIIITDTEISEFDVTKSKNVP
KQASASAENEQVGVFANDNGFEDRTSEQYETASAETASSLVKNAAQSSAEQSANEMASDDNK
INNLLQ
$\varepsilon_{205}$= 415730 $M^{-1}$ $cm^{-1}$

CnA
MGSSHHHHHHSSGLVPRGSHMSEPKAIDPKLSTTDRVVKAVPFPPSHRLTAKEVFDNDGKPR
VDILKAHLMKEGRLEESVALRIITEGASILRQEKNLLDIDAPVTVCGDIHGQFFDLMKLFEV
GGSPANTRYLFLGDYVDRGYFSIECVLYLWALKILYPKTLFLLRGNHECRHLTEYFTFKQEC
KIKYSERVYDACMDAFDCLPLAALMNQQFLCVHGGLSPEINTLDDIRKLDRFKEPPAYGPMC
DILWSDPLEDFGNEKTQEHFTHNTVRGCSYFYSYPAVCEFLQHNNLLSILRAHEAQDAGYRM
YRKSQTTGFPSLITIFSAPNYLDVYNNKAAVLKYENNVMNIRQFNCSPHPYWLPNFMDVFTW
SLPFVGEKVTEMLVNVLNICSDDELGSEEDGFDGATAAARKEVIRNKIRAIGKMARVFSVLR
EESESVLTLKGLTPTGMLPSGVLSGGKQTLQSATVEAIEADEAIKGFSPQHKITSFEEAKGL
DRINERMPPRRDAMPSDANLNSINKALTSETNGTDSNGSNSSNIQ

CnB
MGNEASYPLEMCSHFDADEIKRLGKRFKKLDLDNSGSLSVEEFMSLPELQQNPLVQRVIDIF
DTDGNGEVDFKEFIEGVSQFSVKGDKEQKLRFAFRIYDMDKDGYISNGELFQVLKMMVGNNL
KDTQLQQIVDKTIINADKDGDGRISFEEFCAVVGGLDIHKKMVVDV

CnA/B
$\varepsilon_{280}$= 53290 $M^{-1}$ $cm^{-1}$

Calmodulin
GSADQLTEEQIAEFKEAFSLFDKDGDGTITTKELGTVMRSLGQNPTEAELQDMINEVDADGN
GTIDFPEFLTMMARKMKDTDSEEEIREAFRVFDKDGNGYISAAELRHVMTNLGEKLTDEEVD
EMIREADIDGDGQVNYEEFVQMMTAK
$\varepsilon_{280}$= 2980 $M^{-1}$ $cm^{-1}$

**SUPPLEMENTARY FIGURES ➔**

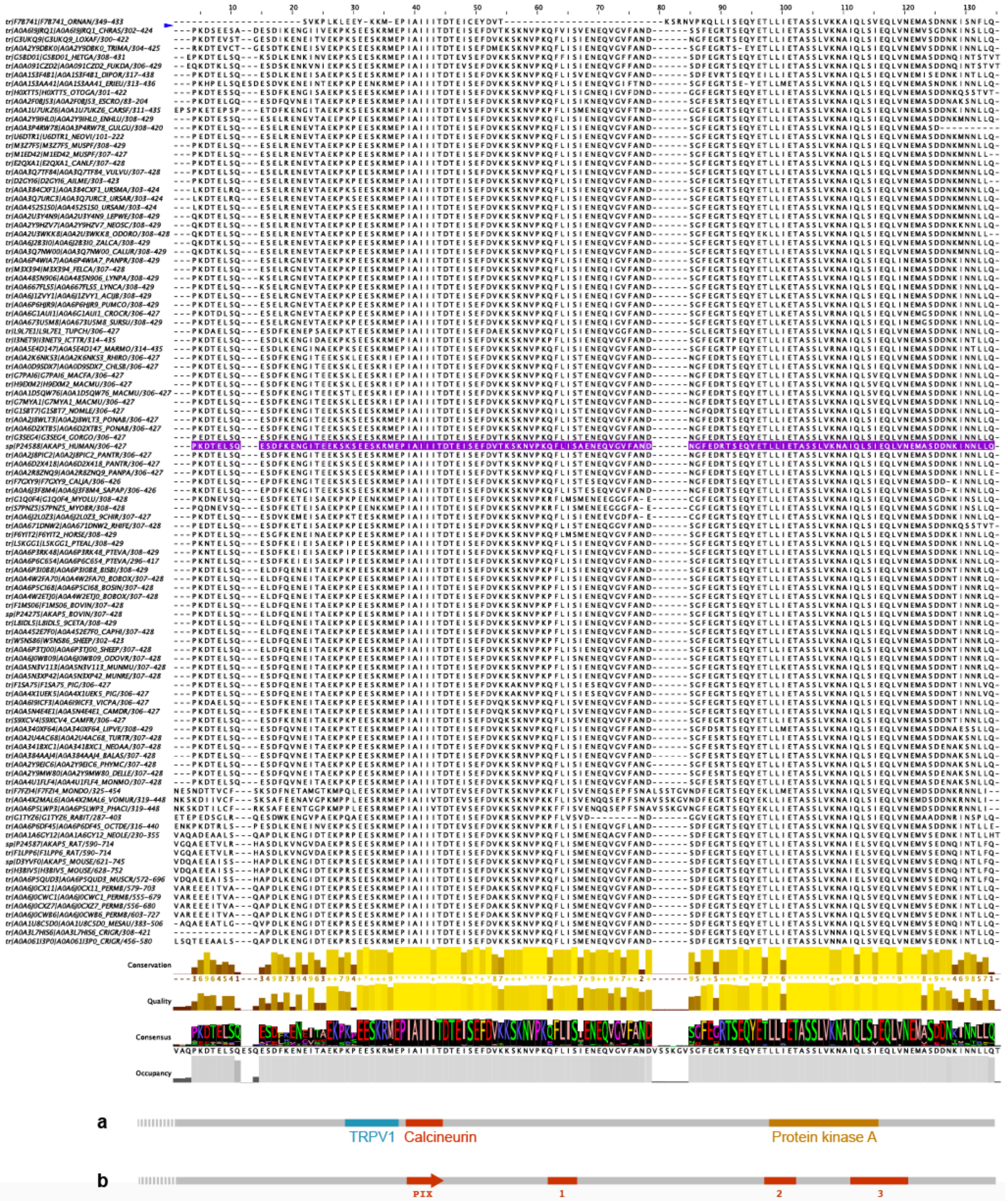


**Supplementary Figure S1: Sequence alignment of the C-terminal region of AKAP5**. All AKAP5 with a PIAIIIT (or similar) SLiM are shown, taken from Uniprot. A region of high conservation runs from residue 306 to 427 in the human sequence (purple highlight). (**a**) Positions of known SLiMs. (**b**) Positions of the three secondary binding sites established in this study, relative to the PxIxIT SLiM. Figure prepared using Jalview v2 (A. M. Waterhouse, J. B. Procter, D. M. A. Martin, M. Clamp, G. J. Barton, Bioinformatics. 25, 1189–1191 (2009)).

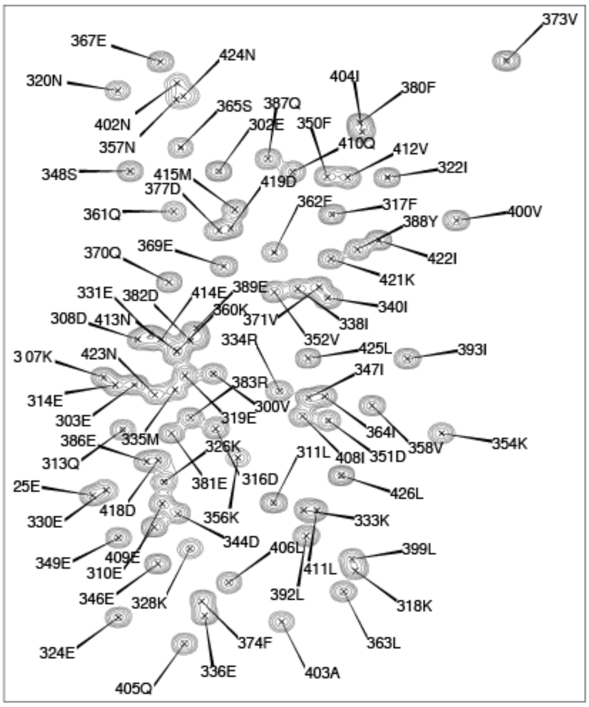


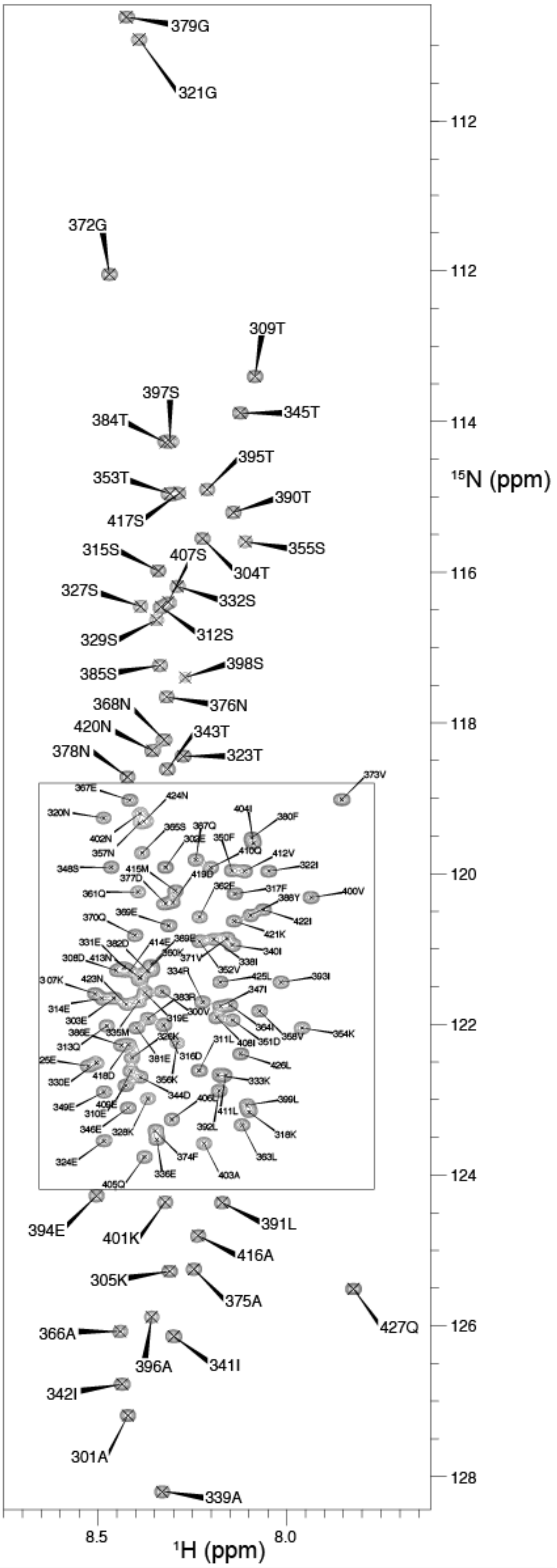


**Supplementary Figure S2: Backbone $^{1}H$,$^{15}N$ chemical shift assignments of AKAP5c**. $^{15}N$-HSQC of cAKAP5 (100 µM in 10 mM PIPES pH 6.5, 150 mM NaCl and 10-15% $^{2}H_2O$). Experiment recorded at 25 °C on Bruker DRX800 spectrometer. Assignments also deposited in the BMRB with accession number XXXX.

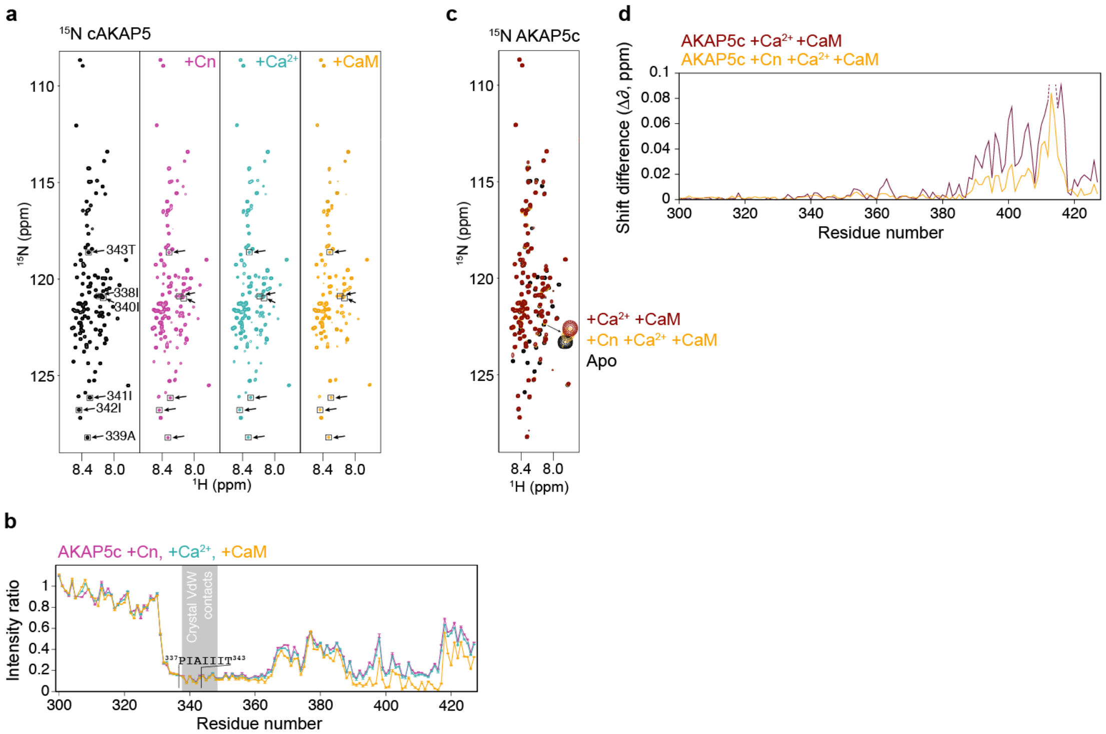


**Supplementary Figure S3: Calmodulin competes with Calcineurin for a distal, non-PxIxIT motif.** (**a**) $^{15}$N-labelled AKAP5c titrated sequentially with Calcineurin (Cn; magenta), $Ca^{2+}$ (cyan) and Calmodulin (CaM; yellow), followed by HSQC. Peaks from non-proline residues in $^{337}$PIAIIIT$^{343}$ are boxed with arrows. No changes were seen on addition of $Ca^{2+}$ to a large excess (20:1). AKAP5c/Calcineurin binding is therefore calcium-independent. However, some non-PIAIIIT peaks further shift and broaden on addition of Calmodulin, mapped in (**b-d**). (**b**) Intensity changes relative to $^{15}$N-AKAP5c alone. PIAIIIT-binding of Calcineurin is unaffected by addition of CaM, but further peak attenuation is observed in the region 390-417. Molar ratios of 1:1 Cn:AKAP, 1:20:1 Cn:$Ca^{2+}$:AKAP and 1:20:1:1 Cn:$Ca^{2+}$:CaM:AKAP are shown. Grey boxed region: van der Waals contacts in the X-ray crystal structure (3LL8). (**c**) Independent binding of CaM/AKAP5c tested by $^{15}$N-labelled AKAP5c (black) titrated with $Ca^{2+}$ and Calmodulin (CaM), followed by HSQC. Yellow: Calcineurin (Cn) present (from **a**), Maroon: Cn absent. Molar ratios 1:20:1:1 Cn:$Ca^{2+}$:CaM:AKAP and 0:20:1:1 Cn:$Ca^{2+}$:CaM:AKAP are shown. Chemical shift changes are seen for 390-417 regardless of the presence of Cn, indicating CaM/AKAP binding is independent of Cn. However, the peaks move further along the shift trajectory in the absence of Cn. (**d**) Chemical shift changes relative to $^{15}$N-cAKAP5 alone. Larger changes are observed in the absence of Cn. Calmodulin therefore appears to compete with Calcineurin for binding to residues 390-417, a region ca. 50 residues from the canonical PxIxIT site that coincides with the PKA SLiM (Fig. 1).

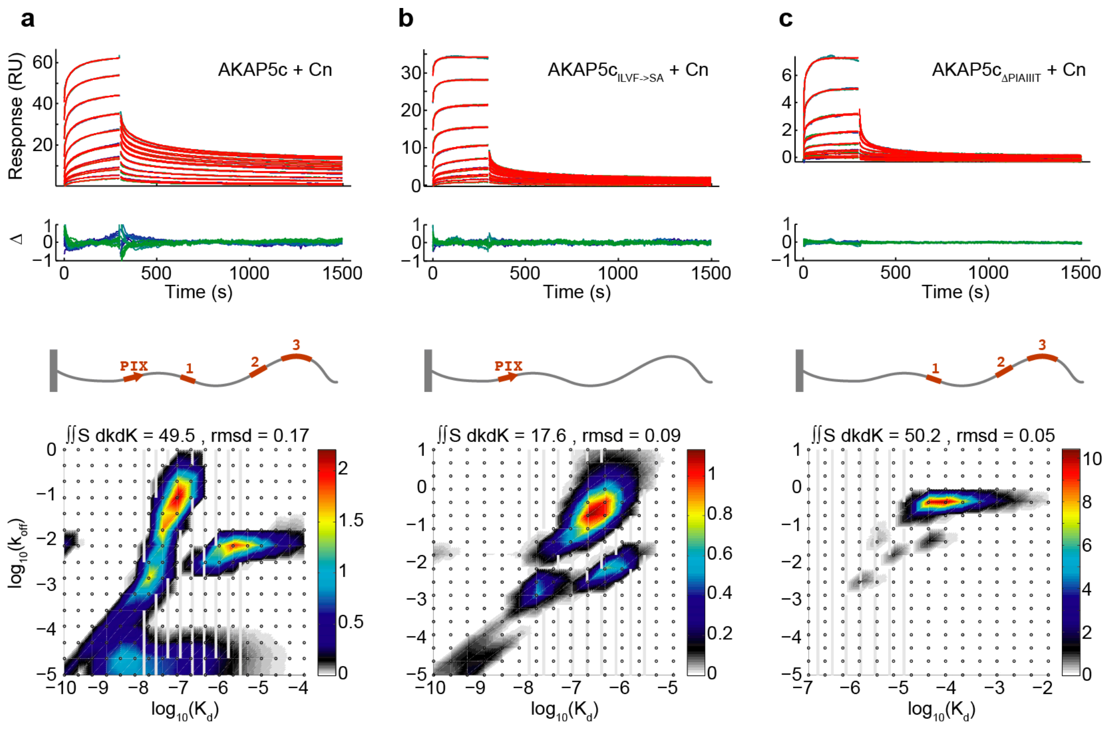


**Supplementary Figure S4: Surface site heterogeneity of AKAPc modelled in EVILFIT (P. Schuck, H. Zhao, *Methods Mol. Biol.* 627, 15–54 (2010)).** (*Top*) SPR response curves of Calcineurin injected over a streptavidin surface pre-immobilised with (**a**) AKAP5c, (**b**) AKAP5c$_{ILVF\text{->}SA}$ or (**c**) AKAP5c$_{\Delta PIAIIIT}$. Raw data repeats in blue and green, fit in red, Δ = residuals to the fit. Calcineurin concentrations were 2-fold dilutions from 2.56 µM (AKAP5c and AKAP5c$_{ILVF\text{->}SA}$) or from 10 µM (AKAP5c$_{\Delta PIAIIIT}$). (*Bottom*) Distribution plots of $k_{off}$ vs $K_d$. The optimal $k_{off}$ and $K_d$ ranges for the fitting were chosen so as to include all the $k_{off}/K_d$ site classes present in each data set, and to achieve the best fit (minimum rmsd). Points were distributed according to the 'Autogrid' function.

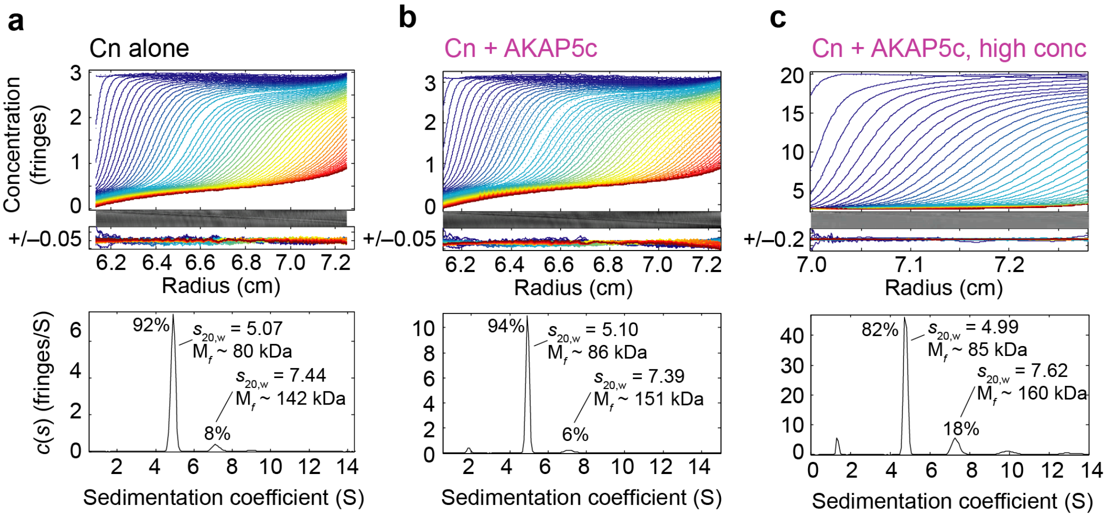


**Supplementary Figure S5:** Sedimentation velocity continuous *c*(*s*) distribution of (a) Calcineurin alone at 7.5 µM concentration in 10 mM Tris/HCl pH 7.4, 150 mM NaCl, 5 mM DTT, (b) as (a) but with added cAKAP5c, and (c) Calcineurin/AKAP5c at high concentration (75 µM each) in the same conditions as for NMR (10 mM PIPES pH 6.5, 150 mM NaCl, no DTT). The average frictional ratio ($f/f_0$) increased from 1.31 to 1.37 in the presence of AKAP5c, indicating that the addition of frictional drag due to protein disorder (that retards sedimentation) counteracts the effect of the additional mass of AKAP5c (14.5 kDa), leading to only a small overall effect on *s* for the complex.

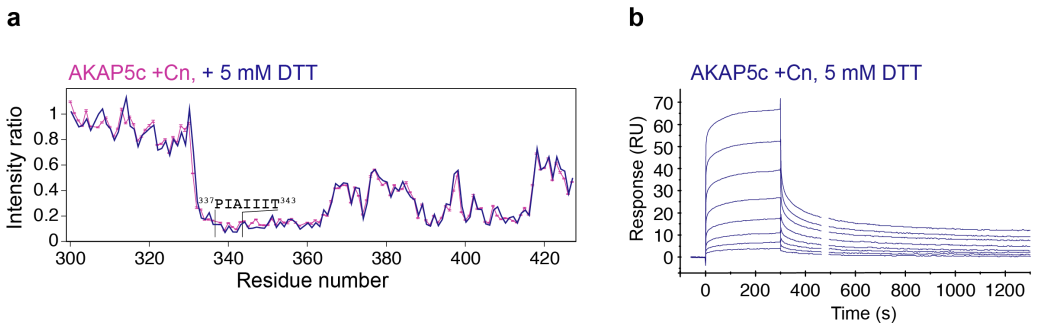


**Supplementary Figure S6: cAKAP/Calcineurin binding by NMR and SPR in strongly reducing conditions. (a)** NMR: $^{15}$N-HSQC intensity changes to AKAP5c on sequential addition of Calcineurin (Cn; magenta) and 5 mM DTT (purple). The proteins are each at 100 µM and were incubated with DTT at 5 mM. (**b**) SPR: Calcineurin injected over a streptavidin surface pre-immobilised with AKAP5c. Calcineurin concentrations were 2-fold dilutions from 2.56 µM. All buffers contained 5 mM DTT and all protein solutions were pre-equilibrated in 5 mM DTT before injection.

**a**

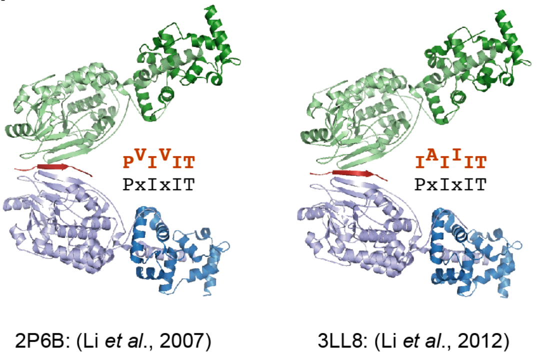


**b**

| | PxIxIT | $K_d$ or $K_i$ (μM) | |
|---|---|---|---|
| | **PPAIIIT** | 0.08 | → PAIIIT / PxIxIT |
| AKAP5 | **PIAIIIT** | 0.4 | |
| | **PPAIIIA** | 12 | |
| | **PIAIIIA** | 39 | |
| | **PVIVIT** | 0.5 | |
| TRESK | **PQIIIS** | 5 | |
| NFAT | **PRIEIT** | 25 | → PRIEIT / PxIxIT |
| Hph1 | **PVIAVN** | 250 | |

**Supplementary Figure S7: Reported crystal structures and affinities of Calcineurin and various PxIxIT peptides.** (**a**) Crystal structures of Calcineurin bound to PVIVIT and PIAIIIT peptides (2P6B [1]; 3LL8 [2]), in which the stoichiometry in the asymmetric unit is 2:1 as a second Calcineurin heterodimer engages with the opposite 'x' side of the PxIxIT motif. (**b**) Affinities of various PxIxIT peptides (taken from [1,2]).

1. H. Li, L. Zhang, A. Rao, S. C. Harrison, P. G. Hogan, *J. Mol. Biol.* **369**, 1296–1306 (2007).
2. H. Li *et al.*, *Nat. Struct. Mol. Biol.* **19**, 337–345 (2012).
